# Modelling cost-effectiveness of tenofovir for prevention of mother to child transmission of hepatitis B virus infection in South Africa

**DOI:** 10.1101/483966

**Authors:** Jolynne Mokaya, Edward Burn, Cynthia Raissa Tamandjou, Dominique Goedhals, Eleanor Barnes, Monique Andersson, Rafael Pinedo-Villanueva, Philippa C Matthews

## Abstract

In light of sustainable development goals for 2030, an important priority for Africa is to have affordable, accessible and sustainable hepatitis B virus (HBV) prevention of mother to child transmission (PMTCT) programmes, delivering screening and treatment for antenatal women and implementing timely administration of HBV vaccine for their babies. We developed a decision-analytic model simulating 10,000 singleton pregnancies to assess the cost-effectiveness of three possible strategies for deployment of tenofovir in pregnancy, in combination with routine infant vaccination: **S1**: no screening nor antiviral therapy; **S2**: screening and antiviral prophylaxis for all women who test HBsAg-positive; **S3**: screening for HBsAg, followed by HBeAg testing and antiviral prophylaxis for women who are HBsAg-positive and HBeAg-positive. Our outcome was cost per infant HBV infection avoided and the analysis followed a healthcare perspective. S1 predicts 45 infants would be HBV-infected at six months of age, compared to 21 and 28 infants in S2 and S3, respectively. Relative to S1, S2 had an incremental cost of $3,940 per infection avoided. S3 led to more infections and higher costs. Given the long-term health burden for individuals and economic burden for society associated with chronic HBV infection, screening pregnant women and providing tenofovir for all who test HBsAg+ may be a cost-effective strategy for South Africa.

## INTRODUCTION

In order to meet targets set by International Sustainable Development Goals for the elimination of Hepatitis B Virus (HBV) infection as a public health problem by the year 2030 (1), enhanced efforts are required to reduce the incidence of new cases. Strategies that set out to achieve this need to be carefully evaluated, both on the grounds of effect on individual and population health, and also based on value for money. HBV infection is highly endemic in many low/middle income settings (2), where economic evaluations are particularly important for informing the appropriate deployment of limited health care resources.

Prevention of mother to child transmission (PMTCT) is a cornerstone of HBV elimination strategies. Reducing vertical transmission is crucial to population health, as up to 90% of neonates who are exposed to HBV perinatally become chronic carriers, compared to only 5% of those exposed as adults (3,4). Current recommended practice includes screening antenatal women for infection using hepatitis B surface antigen (HBsAg), and risk-stratification based on further laboratory tests for hepatitis B e-Antigen (HBeAg) and/or HBV DNA viral load (VL), which can be used to stratify the risk of vertical transmission. PMTCT guidelines suggest administering a three dose HBV vaccine to all infants with the first dose administered within 24 hours of birth and two other doses provided at 6 and 10 weeks respectively, as well as administering hepatitis B immunoglobulin (HBIg) to high risk babies in the first day of life (5,6). Together, these strategies reduce the rate of vertical transmission by 85%-95% (7), representing a crucial component of efforts towards HBV elimination (8–10). However, breakthrough transmission can occur, especially among mothers with high HBV VL (11), in settings where the first dose of vaccine is delayed, and where HBIg is not available. The use of antivirals during pregnancy can therefore be included as an additional measure to decrease transmission risk (12). Tenofovir disoproxil fumarate (TDF) has a track record of safety and efficacy in this setting (13,14), and is included in some guidelines for HBV PMTCT (6,15).

HBV is endemic in Africa, with a prevalence of >8% in many populations (16,17). The risk of MTCT in this continent is enhanced by lack of routine antenatal screening (18,19), deliveries taking place outside healthcare facilities, delayed first dose of HBV vaccine until age six weeks by many vaccine programmes, and limited access to TDF and HBIg (16,18). A meta-analysis of data from sub-Saharan Africa (sSA) demonstrated a perinatal transmission rate of 38% among women who tested positive for HBsAg and HBeAg, in the absence of any PMTCT interventions (20). An important priority for Africa is to have affordable, accessible and sustainable PMTCT programmes that deliver screening and treatment for antenatal mothers, and oversee timely administration of HBV birth vaccine for their babies (8,21). Providing antiviral treatment for HBV PMTCT relies on HBV diagnosis, but currently there is not widespread access to laboratory-based assays for HBsAg, HBeAg and HBV DNA viral load, and many antenatal programmes do not provide routine HBV screening.

In order to identify the most cost-effective approach to HBV PMTCT we have evaluated antenatal screening for HBV infection using standard laboratory assays for HBsAg and treating HBsAg-positive women with TDF, based on South Africa as a model situation. In recognising the important role that lateral flow assays can play in point of care testing (POCT) for HBsAg, we also assessed the cost effectiveness of this approach to diagnostic screening (19,22). In South Africa, the majority of pregnant women are not routinely screened for HBV infection, vaccination begins at six weeks of age and HBIg is not available. Modelling the cost-effectiveness of HBV PMTCT interventions allows for a combined analysis of clinical outcomes and potential budget impact, which is fundamental to inform financial investment in HBV elimination programmes in sSA countries.

## RESULTS

### Cost-effectiveness of strategies for HBV PMTCT under base case assumptions

We modelled three different approaches to peri-partum TDF prophylaxis: **S1**: no screening or antiviral therapy; **S2**: screening and antiviral prophylaxis for all women who test HBsAg-positive; **S3**: screening for HBsAg, followed by HBeAg testing and antiviral prophylaxis for women who are HBsAg-positive and HBeAg-positive (Fig.1). Based on the simulated scenario of 10,000 singleton pregnancies in South Africa, with no antiviral therapy for HBV PMTCT (S1), and using a laboratory-based assay for HBsAg detection, our model predicts 45 infants would be infected with HBV at age six months (incidence 0.45%). If antiviral prophylaxis interventions are employed based on S2 and S3, this would be reduced to 21 and 28 infants, (incidence 0.21% and 0.28%) respectively, at the cost of deploying the intervention for the whole population. Compared to S1, cost per infection avoided was $3,940 for S2, lower than for S3 which would be more costly and avoid a lower number of infections (Table 1; Fig.2).

**Fig. 1:**
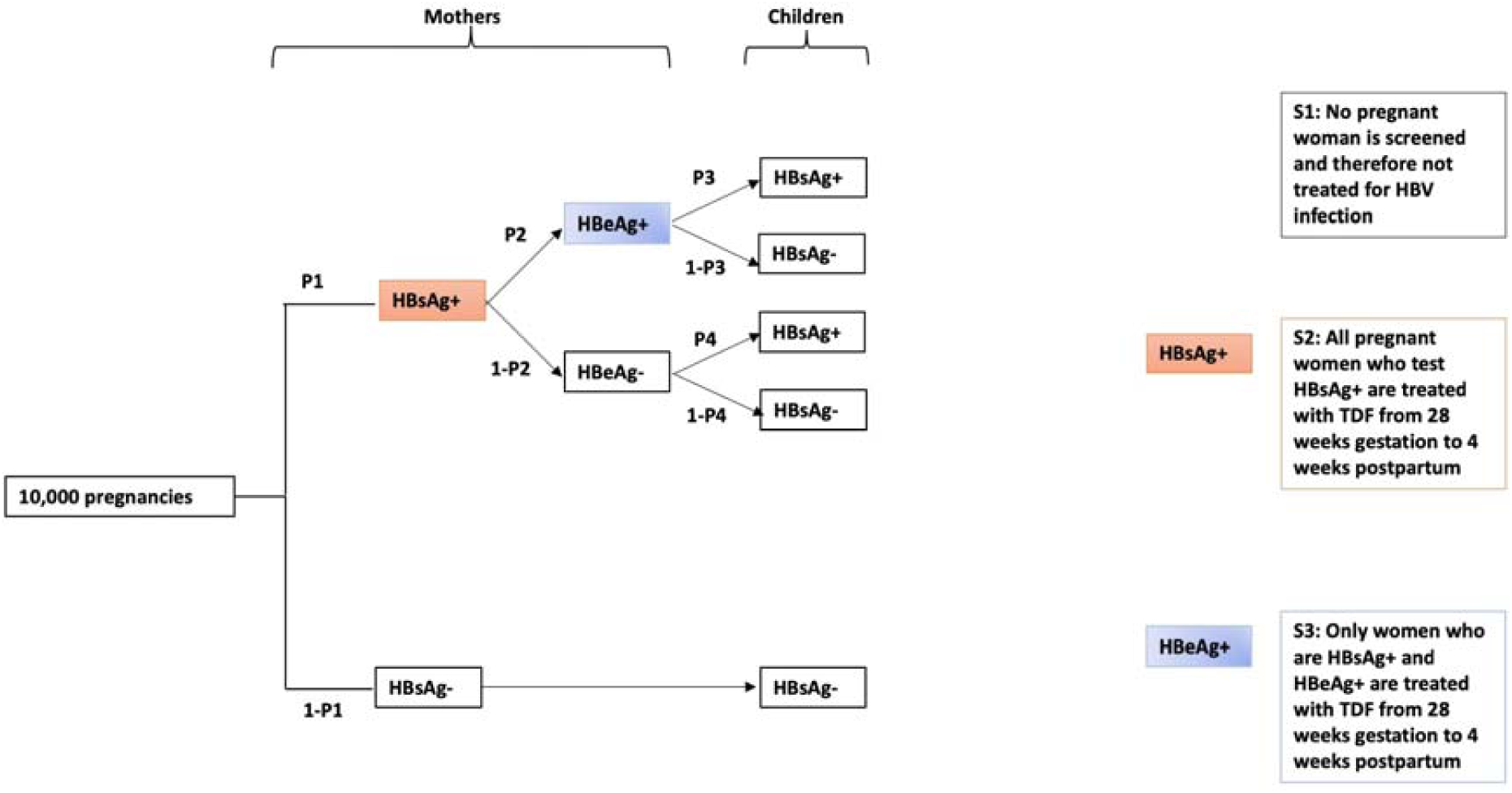
Decision tree showing possible approaches to evaluation of HBV infection in a simulated cohort of 10,000 antenatal women, and in their infants at age six months. P1 is the probability of women testing positive for HBsAg; P2 is the probability of women who are positive for HBsAg also testing positive for HBeAg; P3 is the probability of infants born to HBsAg+/HBeAg+ mothers testing HBsAg+ at 6 months of age; P4 is the probability of infants born to HBsAg+/HBeAg-mothers testing HBsAg+ at 6 months of age. HBsAg: Hepatitis B surface antigen; HBeAg: Hepatitis B e antigen; TDF: Tenofovir disoproxil fumarate.

**Table 1:** Cost-effectiveness results under base case assumptions for screening for HBsAg using laboratory-based assay on a simulated birth cohort of 10,000 live singleton infants.

| Strategy | Number of infant HBV infections<sup>a</sup> (95% CI) | Cost of deploying the intervention for the whole population<sup>b</sup> (n=10,000) in \$ (95% CI) | Incremental cost per infection avoided (\$) |
| --- | --- | --- | --- |
| S1 | 45 (29 - 121) | 0 | - |
|  | 21 (14 – 69) | 94,571 | 3,940 |
| S2 |  | (94,487 - 95,509) | (compared to strategy S1) |
| S3 | 28 (19 - 76) | 95,097<br>(94,980 - 97,244) | Dominated<sup>c</sup> (by strategy S2) |

<sup>a</sup> World Health Organisation (WHO) criteria for HBV elimination states an aim of 90% reduction in new chronic infection (1).
<sup>b</sup> Price of TDF estimated at \$2.48/month for strategies S2 and S3 (38).
<sup>c</sup> S3 is dominated due to both higher costs and higher infections compared to S2.
CI = confidence interval.

**Fig. 2:**
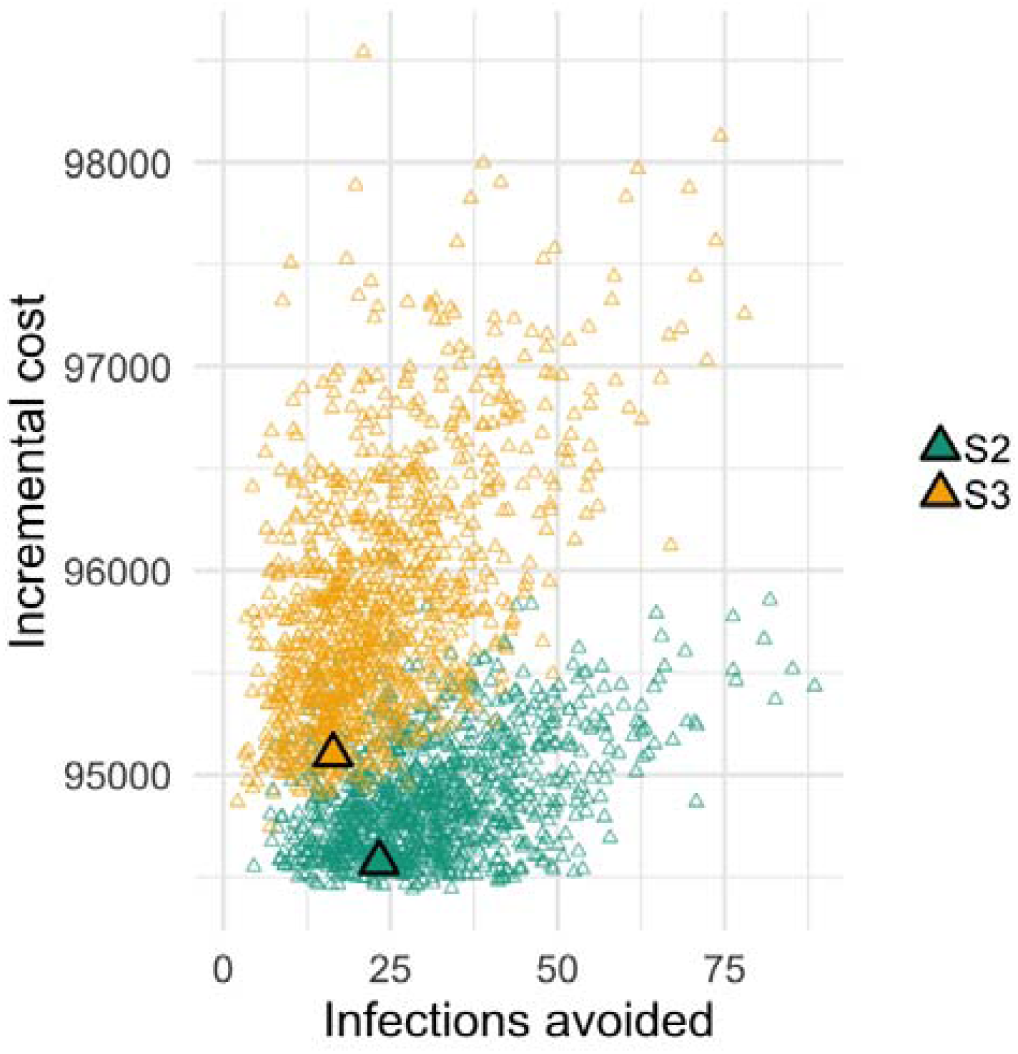
Cost-effectiveness (CE) plane for HBsAg using laboratory assay in a simulated cohort of 10,000 antenatal women. **S2**: All pregnant women are screened for HBsAg; those who test positive are treated with TDF from 28 weeks’ gestation to 4 weeks post-partum. **S3:** All pregnant women are screened for HBsAg, those who test positive are screened for HBeAg. Only those who are HBeAg positive are treated with TDF from 28 weeks’ gestation to 4 weeks post-partum.

### Cost-effectiveness of strategies for HBV PMTCT under scenario analyses

#### (i) Antenatal screening for HBV infection using a rapid point of care test (POCT)

POCT for HBsAg was cheaper in comparison to laboratory-based HBsAg assay (for S2: POCT $23,401 vs lab test $94,571 and for S3: POCT $23,979 vs lab test $95,097); (Table 1 & Table 2). Based on reasonable assumptions about sensitivity of HBsAg detection via laboratory ELISA vs POCT (100% and 97.6%, respectively(19)), from our simulated cohort, laboratory testing would detect all maternal HBV infections (n=360 in our hypothetical cohort of 10,000), while the POCT approach would detect 351 and miss nine cases of infection. This loss of sensitivity translates into a marginal decrease in cost-effectiveness: the incremental cost per infection avoided is estimated at $1,017 compared to $978 if the test had 100% sensitivity.

**Table 2.** Cost-effectiveness results of different PMTCT strategies under scenario analyses on a simulated birth cohort of 10,000 live singleton infants.

| Strategy | Number of infant HBV infections<sup>a</sup> (95% CI) | Cost of deploying the intervention for the whole population (n=10,000) in \$ (95% CI) | Incremental cost per infection avoided (\$) |
| --- | --- | --- | --- |
| Scenario analysis (i): POCT testing |  |  |  |
| S1 | 45 (28 – 119) | 0 |  |
|  | 22 (14 – 73) | 23,401 | 1,017 |
| S2<sup>b</sup> |  | (23,330 – 24,243) | (compared to strategy S1) |
| S3<sup>b</sup> | 28 (19 – 78) | 23,979<br>(23,882 – 25,937) | Dominated<sup>c</sup><br>(by strategy S2) |
| Scenario analysis (ii): Universal birth dose vaccination |  |  |  |
| S1 | 45 (28 - 123) | 0 | - |
| S2<sup>d</sup> | 15 (11 - 82) | 94,571<br>(94,489 - 95,503) | 3,152 (compared to<br>strategy S1) |
| S3<sup>d</sup> | 24 (17 - 84) | 95, 097<br>(94,981 – 97,291) | Dominated<sup>c</sup> (<br>by strategy S2) |
| Scenario analysis (iii): TDF resistance |  |  |  |
| S1 | 45 (29 – 119) | 0 | - |
| S2<sup>d</sup> | 21 (14 – 71) | 94,571<br>(94,496 – 95,515) | 3,940 (compared to<br>strategy S1) |
| S3<sup>d</sup> | 28 (18 – 79) | 95,097<br>(94,980 – 97,252) | Dominated<sup>c</sup> (<br>by strategy S2) |

<sup>a</sup> World Health Organisation (WHO) criteria for HBV elimination states an aim of 90% reduction in new chronic infection (1).
<sup>b</sup> Price of TDF estimated at \$2.42/month for strategies S2 and S3 when using POCT.
<sup>c</sup> S3 is dominated due to both higher costs and higher infections compared to S2.
<sup>d</sup> Price of TDF estimated at \$2.48/month for strategies S2 and S3 (38).
CI = confidence interval.

#### (ii) Recommended PMTCT guidelines: Universal birth dose vaccination and HBIg

Incorporating universal birth dose vaccination and HBIg into our model, reduced the number of infants infected at six months and cost per infection avoided (S2: 15 infants infected; S3: 24 infants infected; and cost per infection avoided: $3,152), (Table 2), compared to when PMTCT intervention includes only TDF and HBV vaccination beginning at six weeks, (S2: 21 infants infected; S3: 28 infants infected; and cost per infection avoided: $3,940) (Table 1).

#### (iii) HBV resistance to TDF

When we accounted for potential HBV resistance to TDF, based on a prevalence of drug resistance estimated at 0.8% (23), there was no change in incremental cost per infection avoided in S2.

## Data visualisation

We reproduced our decision model as an R Shiny application (source code available here: https://github.com/edward-burn/PMTCT-HBV-cost-effectiveness-analysis. This can be used to estimate cost-effectiveness of our three strategies using different input parameters. This will allow the analysis to be re-run for the same comparison of strategies but in contexts where the epidemiology of HBV infection, and costs of interventions, are different.

## DISCUSSION

Antenatal screening and treating of pregnant women for HBV is not routinely performed in many settings in sSA, despite the high population prevalence of HBV (19,24–27). In resource-limited settings, screening and providing prophylactic TDF for all pregnant women who are HBsAg positive may be the most efficient strategy, especially when using a POCT. These results are derived from a theoretical model, and careful consideration is needed for application to different real world settings.

Based on existing data for South Africa, screening and treating women with TDF from 28 weeks of pregnancy to four weeks post-delivery reduces the number of infants with HBV at six months. Previous studies have also reported on the safety and effectiveness of TDF for HBV PMTCT (13,28). There is evidence for the cost-effectiveness of combining antiviral therapy during pregnancy with HBV immunoprophylaxis, as highlighted by a systematic review and meta-analysis carried out in China (29) and a study conducted in North America (30). Interestingly, a recent study in Thailand conflicted with these findings by reporting no significant benefit of maternal TDF for PMTCT (31). The lack of benefit from TDF in this case may be because all infants in the Thai study received birth dose of both HBIg and HBV vaccine, followed by four additional doses of HBV vaccine; this enhanced immunoprophylaxis probably accounts for the low rate of HBV transmission even among mothers who did not receive TDF. However, it is difficult to apply any of these findings to sSA, which differs in having limited (or no) access to HBIg, gaps in vaccine coverage, and delays in the first dose of HBV vaccine (frequently until age six weeks) (16,18).

In this study, we provide a head-to-head economic analysis of simulated screening for HBsAg alone vs. use of HBsAg in combination with risk stratification using HBeAg. Limited resources and infrastructure have impeded antenatal diagnosis and treatment of HBV (27), and stratification of HBsAg-positive antenatal women with HBV DNA level and/or HBeAg to determine eligibility for TDF incurs further cost. Given that TDF is becoming more accessible for HBV treatment in Africa (10,32), and has a well-established safety record as a result of antenatal use in HIV infection, treating all HBsAg positive women should be safe and practical, as well as cost-effective. As this option is simple to implement, it is therefore also most likely to be successful.

Given the high prevalence of HBV in Africa (27), POCT is an appealing route to increasing diagnosis and therefore access to treatment. Although POCT proves advantageous on cost grounds, it is less sensitive than laboratory assays. Furthermore, laboratory support remains key for monitoring response to treatment, identifying and monitoring drug resistance, and evaluating prognosis.

There is a potential risk that increasing population exposure to TDF may increase selection of resistance (32). However, the genetic barrier to TDF resistance is high, and more data are needed to determine the prevalence and clinical significance of putative drug resistance mutations. When accounting for an estimated background rate of HBV resistance to TDF, we show no change in the cost per infection avoided. While we recognise that TDF resistance is not currently a significant clinical concern, the modelling approach we have developed allows drug resistance to be factored in.

Despite the potential for increased risk of HBV MTCT in HIV/HBV coinfection (33), there is evidence to show that HIV has very little effect on HBV interventions (34). Since HIV guidelines now recommend commencement of antiretroviral treatment as soon as HIV diagnosis is made (35), the impact of HIV on HBV PMTCT is further reduced.

We used a simple model with only a single outcome measure, thus overlooking other possible risks/benefits of antiviral therapy, including side-effects, drug interactions, and rebound hepatitis after treatment cessation. However, extensive experience has led to the inclusion of TDF in first-line regimens for HIV, including in pregnant women. Our base values were obtained from published literature, but there are limited data for HBV epidemiology and transmission for most African populations; we have acknowledged the lack of certainty around certain parameters by including confidence intervals. HBV DNA testing is the gold standard approach to HBV diagnosis, but is often not available in resource-limited settings; we therefore used HBeAg status as a surrogate marker to represent infectivity and high VL and assumed 100% sensitivity for laboratory-based assays for HBsAg and HBeAg. This does not account for occult infection and other false negatives and may lead to an over-estimation of the cost-effectiveness of maternal TDF.

The assumption that the first dose of HBV vaccine is delayed until age six weeks is an over-simplification: if birth dose vaccine is given more widely or targeted for infants born to HBsAg-positive mothers, more new infections will be averted.

We used data for South Africa, but recognise that HBV prevalence is substantially higher in other sSA settings (frequently reported at ≥8%, the WHO threshold for high endemicity (16,17)) and this may influence the cost-effectiveness of proposed interventions. The low prevalence rate that we derived from South Africa may be a reflection that many antenatal women themselves received HBV vaccination as infants (if born after 1995). To facilitate cost-effectiveness simulations in other settings where antenatal HBV prevalence might be higher, we have provided an on-line interactive tool.

The findings of this study are applicable to settings where the cost of screening and treatment is publicly funded; in situations where individuals meet the cost of their own screening and treatment, our results cannot be applied to informing public health strategy. Our analysis did not consider the need for healthcare and laboratory infrastructure to support our proposed interventions, costs of monitoring pregnant women during TDF treatment, costs for clinical visits and counselling for mothers testing HBsAg positive, leading to an underestimation of the total costs included in providing robust PMTCT. However, this is offset by the potentially very substantial lifelong costs of chronic HBV infection, both to individuals and to society, arising from morbidity and mortality typically affecting young and middle-aged adults. This is difficult to quantify but leads to a substantial burden of chronic disease with economic consequences for the health-care system, as well as imposing financial consequences on families and society (21,36).

## Conclusion

We have developed a simple, theoretical model that allows us to estimate the impact of providing TDF to antenatal women in a lower/middle income country setting, either based on HBsAg status alone, or incorporating risk-stratification with HBeAg. These strategies reflect safe, practical interventions that could reasonably be deployed in many settings. There remains an urgent need for more data to underpin the relevant epidemiology, risks of MTCT, and relative benefits of different interventions in settings across sSA. In order to drive progress towards 2030 elimination targets, sustained investment is required to drive improvements in clinical services, provide universal access to antenatal screening, improve education of the public and health-care workers, and underpin robust deployment of PMTCT interventions including vaccination and TDF therapy.

## METHODS

### Target population and study perspective

We used a hypothetical cohort of 10,000 pregnant South African women to evaluate the cost-effectiveness of three different strategies for antiviral therapy with TDF:

#### Strategy 1 (S1): No TDF prophylaxis

No pregnant woman is screened for HBsAg and therefore no HBV treatment is given perinatally. To date, this is the situation in many resource-limited settings in sSA.

#### Strategy 2 (S2): Screening and TDF prophylaxis for all women who test HBsAg positive

All pregnant women are screened for HBsAg; those who test positive are treated with TDF from 28 weeks’ gestation to four weeks post-partum.

#### Strategy 3 (S3): Screening and TDF prophylaxis for women who are both HBsAg positive and HBeAg positive

All pregnant women are screened for HBsAg; those who test positive are then screened for HBeAg. Only those who are HBeAg positive are treated with TDF from 28 weeks’ gestation to four weeks post-partum.

We populated our model with data using a healthcare system perspective and therefore only considered costs that would be directly incurred by the Department of Health (DOH). These costs relate to screening and treatment of HBV infection during pregnancy. Individual patient information was not included in this analysis. Ethics approval was not required for this study.

### Assumptions

For the purposes of this analysis, we made a number of modelling assumptions:

- All pregnancies are singleton and result in the delivery of a live infant;
- All infants receive HBV vaccine at 6, 10 and 14 weeks, as per current standard practice in South Africa (19). Even for babies born to mothers who are HBsAg+, birth dose vaccination is not offered within 24 hours (due to lack of maternal screening);
- HBV screening uptake is 100% among antenatal women;
- HBsAg and HBeAg are tested using laboratory assays which are 100% sensitive and specific;
- Infants born to mothers who were HBsAg negative were assumed also to be HBsAg negative;
- We made assumptions around our point estimates due to lack of data.

### Theoretic modelling approach

A decision tree providing a framework for comparison of the three strategies is shown in Fig. 1. The decision tree considers the period covering weeks 28 through 40 of gestation up to six months postpartum. Model structure was the same for all strategies. HBsAg-positive mothers had a defined probability of being HBeAg positive or negative; these probabilities were the same for all strategies. Children whose mothers were HBsAg positive had a probability of being HBsAg positive themselves (i.e. a case of MTCT). This probability depended on whether the mother was also HBeAg positive and whether antiviral prophylaxis was received by the mother during pregnancy, which varied between strategies.

### Measurement of effectiveness

Table 3 shows parameters used for input in our model. We searched the published literature, and identified four studies from within the last six years that reported the prevalence of HBV infection among pregnant women in South Africa (19,24–26). To calculate prevalence of HBV infection, we combined the total number of women from each study who tested HBsAg-positive and divided these by the total number of individuals included in these cohorts. From these pooled data, the overall prevalence of HBsAg among pregnant women was 3.6% (129/3614). HBeAg status was reported for 126 of these 129; the prevalence of HBeAg among HBsAg+ women was 29/126 (23%).

**Table 3:**
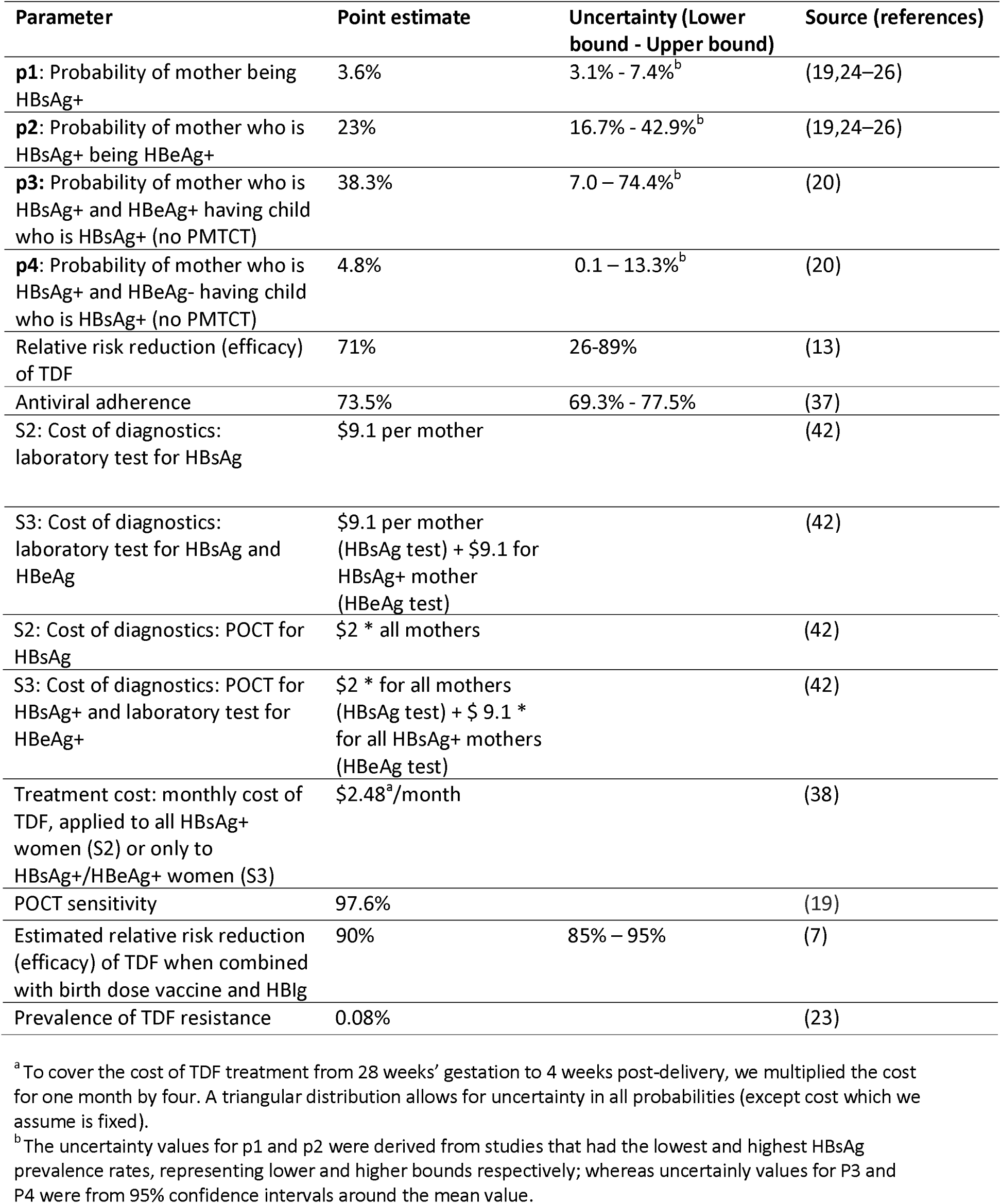
Parameters used for input into a model of cost-effectiveness of tenofovir for PMTCT in South Africa, based on a simulated cohort of 10,000 antenatal women and their babies

We derived point estimates for perinatal transmission with and without antiviral therapy during pregnancy from a published systematic review and a meta-analysis (13,20). In the absence of intervention (HBV vaccination, HBIg and/or antiviral therapy), the perinatal transmission rate for HBsAg+/HBeAg+ mothers in sSA was estimated at 38%, and for HBsAg+/HBeAg-mothers was 4.8% (20). Due to lack of data, we used these estimates in our model, although we assumed that infants were vaccinated starting at the age of six weeks.

There is only one randomised control trial that examines the efficacy of TDF on HBV PMTCT, in which perinatal TDF reduced the risk of infant HBsAg seropositivity by 71% (12). However, data from a systematic review and meta-analysis estimate this figure as 77% (13). We therefore reduced the perinatal transmission rate for women receiving no TDF by 71% to obtain a point estimate for perinatal transmission rate for those receiving TDF peripartum prophylaxis; these were estimated at 11.1% for HBeAg-positive women and 1.4% for HBeAg-negative women. We accounted for uncertainty around point estimates, as the original study was conducted in China and with a different protocol (combining TDF during pregnancy in combination with birth dose vaccine and HBIg).

Due to lack of data on HBV antiviral adherence rates among pregnant women in sSA, we used HIV antiretroviral adherence rate among women during and after pregnancy, obtained from a published systematic literature review which included studies from low-income, middle-income, and high-income countries (37); this is estimated at 73.5%. We incorporated an expected adherence of 73.5% into the model, assuming that those mothers who didn’t adhere would incur the full cost of treatment but would have the same probability of transmission as un-treated women.

### Outcomes

Our outcome was the cost per HBV infection avoided among infants at the age of six months. This outcome was selected because it is quantifiable, clinically relevant and could be easily compared between strategies.

### Estimating costs

We derived costs for laboratory assays for HBsAg and HBeAg, and POCT, from a 2015 price list produced by National Health Laboratory Services (NHLS) and DOH, South Africa. We used the cost of TDF from International Medical products price guide for year 2015 (38). We converted the price of HBsAg laboratory assay (R 108.86), HBsAg POCT (R18), HBeAg laboratory assay (R108.86) and HBV DNA test (R1173.32) from South African Rand to USD; 1 Rand = 0.083 USD as at 1^st^ May 2015. HBsAg screening cost applied to all women in S2 and S3; HBeAg screening costs applied to women who tested positive for HBsAg in S3. TDF costs applied to all women who tested positive for HBsAg in S2, and women who tested positive for both HBsAg and HBeAg in S3.

### Scenario analysis

We assessed the impact of three additional scenarios, estimating the number of MTCT cases, costs and incremental cost per infection avoided in each case:

#### (i) Antenatal screening for HBV infection using a rapid point of care test (POCT)

We estimated the cost-effectiveness of using a POCT for HBsAg screening, based on a POCT sensitivity of 97.6% and specificity of 100%, compared to laboratory ELISA HBsAg testing as the gold standard (19). We incorporated this into the model by reassigning the proportion of women expected to test falsely HBsAg-using a POCT into the untreated group and removing the cost of TDF treatment for these individuals.

#### (ii) Recommended PMTCT guidelines: Universal birth dose vaccination and HBIg

To assess the cost-effectiveness of antenatal screening and treating of HBsAg-positive women with TDF in combination with universal birth dose vaccine and HBIg, we estimated that perinatal transmission rate will be reduced by 90% (7). To incorporate the impact of TDF in our cohort, we therefore reduced the perinatal transmission rate by 90%; MTCT rates were thus estimated for HBsAg+ women who are HBeAg+ and HBeAg-at 3.8% and 0.48% respectively.

#### (iii) HBV resistance to TDF

In order to consider the possible impact of drug resistance on the PMTCT strategies outlined here, we estimated the prevalence of TDF resistance based on HBV sequences retrieved from the HBV database (https://hbvdb.ibcp.fr/HBVdb/) (23) accessed on the 18^th^ April 2018. The numerator was the total number of sequences with mutations associated with reduced sensitivity/resistance to TDF (rtA194T, rtN236T, rtS78T),(39,40) and the denominator was the total number of sequences available in the database (49/6287, 0.8%). This prevalence was incorporated into the model by re-assigning 0.8% of those receiving TDF into the untreated category (as although they receive the drug, resistance would render this functionally equivalent to no treatment).

### Analytic methods and definitions

We used the following definitions:

- A strategy was ‘dominated’ if it had a higher expected cost and more predicted cases of HBV MTCT compared to an alternative strategy.
- The incremental cost per HBV infection avoided was determined by dividing the difference in cost between two strategies over the difference in number of infections (41).

We compared the strategies by first applying decision rules to eliminate any strategies which were dominated by others, and then estimated the incremental cost per infection avoided for the remaining strategies.

The combined effect of uncertainty in probability estimates was assessed using probabilistic sensitivity analysis, with 1000 Monte Carlo simulations run. We used triangular distributions to approximate Beta distributions for probabilities. Triangular distributions required a lower and upper bound, with the peak taken as the expected (mean) value. The impact of uncertainty is shown on a cost-effectiveness plane, where sets of incremental costs and number of infections avoided are plotted for S2 and S3, relative to S1.

### Standardised criteria

Our analysis conforms to the standardised criteria for economic evaluations of health interventions (our CHEERS checklist can be reviewed as Supplementary Table S1 on-line:https://doi.org/10.6084/m9.figshare.7265582.v1

## Supporting information

## AUTHORS’ CONTRIBUTIONS

- Conceived the study: JM, MA, PCM, CRT
- Literature review: JM
- Assimilated data to feed into the model : JM, DG, PCM
- Analysed the data : JM, EB, RPV, PCM
- Wrote the manuscript: JM, EB, PCM
- Edited and approved the final manuscript : all authors

## Abbreviations

HBIg: hepatitis B immunoglobulin
HBV: hepatitis B Virus
HBeAg: hepatitis B e antigen
HBsAg: hepatitis B surface antigen
MTCT: mother to child transmission
PMTCT: prevention of mother to child transmission
POCT: point of care test
sSA: sub Saharan Africa
TDF: Tenofovir disoproxil fumarate
VL: viral load

