## Supplementary material for "Modelling cost-effectiveness of tenofovir for prevention of mother to child transmission of hepatitis B virus infection in South Africa"

**CHEERS checklist— Cost-effectiveness of alternative strategies for prevention of**

| **Section/item** | **Item No** | **Recommendation** | **Reported on page No/ line No** |
| --- | --- | --- | --- |
| **Title and abstract** | | | |
| Title | 1 | Identify the study as an economic evaluation or use more specific terms such as “cost-effectiveness analysis”, and describe the interventions compared. | Pg.i: Modelling cost-effectiveness of tenofovir for  prevention of mother to child transmission  of hepatitis B virus infection in South Africa |
| Abstract | 2 | Provide a structured summary of objectives, perspective, setting, methods (including study design and inputs), results (including base case and uncertainty analyses), and conclusions. | Abstract |
| **Introduction** | | | |
| Background and objectives | 3 | Provide an explicit statement of the broader context for the study. | Described in introduction |
|  |  | Present the study question and its relevance for health policy or practice decisions. | Described in introduction |
| **Methods** | | | |
| Target population and subgroups | 4 | Describe characteristics of the base case population and subgroups analysed, including why they were chosen. | Described in methods section: Target population and study perspective |
| Setting and location | 5 | State relevant aspects of the system(s) in which the decision(s) need(s) to be made. | Described in methods section: Target population and study perspective |
| Study perspective | 6 | Describe the perspective of the study and relate this to the costs being evaluated. | Described in methods section: Target population and study perspective |
| Comparators | 7 | Describe the interventions or strategies being compared and state why they were chosen. | Described in methods section: Strategies |
| Time horizon | 8 | State the time horizon(s) over which costs and consequences are being evaluated and say why appropriate. | Described in methods section |
| Discount rate | 9 | Report the choice of discount rate(s) used for costs and outcomes and say why appropriate. | N/A |
| Choice of health outcomes | 10 | Describe what outcomes were used as the measure(s) of benefit in the evaluation and their relevance for the type of analysis performed. | Described in methods section: Outcomes |
| Measurement of effectiveness | 11a | *Single study-based estimates:* Describe fully the design features of the single effectiveness study and why the single study was a sufficient source of clinical effectiveness data. | Described in methods section: Measurement of effectiveness |
|  | 11b | *Synthesis-based estimates*: Describe fully the methods used for identification of included studies and synthesis of clinical effectiveness data. | Described in methods section: Measurement of effectiveness |
| Measurement and valuation of preference based outcomes | 12 | If applicable, describe the population and methods used to elicit preferences for outcomes. | Described in methods section: Measurement of effectiveness |
| Estimating resources and costs | 13a | *Single study-based economic evaluation:*Describe approaches used to estimate resource use associated with the alternative interventions. Describe primary or secondary research methods for valuing each resource item in terms of its unit cost. Describe any adjustments made to approximate to opportunity costs. | Described in methods section: Estimating costs |
|  | 13b | *Model-based economic evaluation:* Describe approaches and data sources used to estimate resource use associated with model health states. Describe primary or secondary research methods for valuing each resource item in terms of its unit cost. Describe any adjustments made to approximate to opportunity costs. | Described in methods section: Estimating costs |
| Currency, price date, and conversion | 14 | Report the dates of the estimated resource quantities and unit costs. Describe methods for adjusting estimated unit costs to the year of reported costs if necessary. Describe methods for converting costs into a common currency base and the exchange rate. | Described in methods section: Estimating costs |
| Choice of model | 15 | Describe and give reasons for the specific type of decision-analytical model used. Providing a figure to show model structure is strongly recommended. | Described in methods section: Theoretic modelling approach |
| Assumptions | 16 | Describe all structural or other assumptions underpinning the decision-analytical model. | Described in methods section: Assumptions |
| Analytical methods | 17 | Describe all analytical methods supporting the evaluation. This could include methods for dealing with skewed, missing, or censored data; extrapolation methods; methods for pooling data; approaches to validate or make adjustments (such as half cycle corrections) to a model; and methods for handling population heterogeneity and uncertainty. | Described in methods section: Analytic methods and definitions |
| **Results** | | | |
| Study parameters | 18 | Report the values, ranges, references, and, if used, probability distributions for all parameters. Report reasons or sources for distributions used to represent uncertainty where appropriate. Providing a table to show the input values is strongly recommended. | Described in methods section: Table 3 |
| Incremental costs and outcomes | 19 | For each intervention, report mean values for the main categories of estimated costs and outcomes of interest, as well as mean differences between the comparator groups. If applicable, report incremental cost-effectiveness ratios. | Described in results section: Table 1 |
| Characterising uncertainty | 20a | *Single study-based economic evaluation:*Describe the effects of sampling uncertainty for the estimated incremental cost and incremental effectiveness parameters, together with the impact of methodological assumptions (such as discount rate, study perspective). | N/A |
|  | 20b | *Model-based economic evaluation:* Describe the effects on the results of uncertainty for all input parameters, and uncertainty related to the structure of the model and assumptions. | Described in results section: Table 1 |
| Characterising heterogeneity | 21 | If applicable, report differences in costs, outcomes, or cost-effectiveness that can be explained by variations between subgroups of patients with different baseline characteristics or other observed variability in effects that are not reducible by more information. | Described in results section: Table 2 |
| **Discussion** | | | |
| Study findings, limitations, generalisability, and current knowledge | 22 | Summarise key study findings and describe how they support the conclusions reached. Discuss limitations and the generalisability of the findings and how the findings fit with current knowledge. | Described in discussion section |
| **Other** | | | |
| Source of funding | 23 | Describe how the study was funded and the role of the funder in the identification, design, conduct, and reporting of the analysis. Describe other non-monetary sources of support. | Financial support |
| Conflicts of interest | 24 | Describe any potential for conflict of interest of study contributors in accordance with journal policy. In the absence of a journal policy, we recommend authors comply with International Committee of Medical Journal Editors recommendations. | Conflict of interest |

For consistency, the CHEERS statement checklist format is based on the format of the CONSORT statement checklist
